# Medical relevance of protein-truncating variants across 337,208 individuals in the UK Biobank study

**DOI:** 10.1101/179762

**Authors:** Christopher DeBoever, Yosuke Tanigawa, Greg McInnes, Adam Lavertu, Chris Chang, Carlos D. Bustamante, Mark J. Daly, Manuel A. Rivas

## Abstract

Protein-truncating variants can have profound effects on gene function and are critical for clinical genome interpretation and generating therapeutic hypotheses, but their relevance to medical phenotypes has not been systematically assessed. We characterized the effect of 18,228 protein-truncating variants across 135 phenotypes from the UK Biobank and found 27 associations between medical phenotypes and protein-truncating variants in genes outside the major histocompatibility complex. We performed phenome-wide analyses and directly measured the effect of homozygous carriers, commonly referred to as “human knockouts,” across medical phenotypes for genes implicated to be protective against disease or associated with at least one phenotype in our study and found several genes with strong pleiotropic or non-additive effects. Our results illustrate the importance of protein-truncating variants in a variety of diseases.

---

Protein-truncating variants (PTVs), genetic variants predicted to shorten the coding sequence of genes, are a promising set of variants for drug discovery since identification of PTVs that protect against human disease provides *in vivo* validation of therapeutic targets^1,2,3,4^. Although tens of thousands of standing germline PTVs have been identified^5,6^, their medical relevance across a broad range of phenotypes has not been characterized. Because most PTVs are present at low frequency, assessing the effects of PTVs requires genotype data from a large number of individuals with linked phenotype data for a variety of diseases and physiological measurements. The recent release of genotype and linked clinical and questionnaire data for 488,377 individuals in the UK Biobank provides an unprecedented opportunity to assess the clinical impact of truncating protein-coding genes at a resolution not previously possible.

## Results

To assess the clinical relevance of PTVs, we cataloged predicted PTVs present in the Affymetrix UK Biobank array and their effects on medical phenotypes from 337,208 unrelated individuals in the UK Biobank study ^7,8^. We defined PTVs as single-nucleotide variants (SNVs) predicted to introduce a premature stop codon or to disrupt a splice site or small insertions or deletions (indels) predicted to disrupt a transcript’s reading frame ^5^. We identified 18,228 predicted PTVs in the UK Biobank array that were polymorphic across 8,750 genes after filtering (Methods, Figure S1). Each participant had 95 predicted PTVs with minor allele frequency (MAF) less than 1% on average, and 778 genes were predicted to be homozygous or compound heterozygous for PTVs with MAF less than 1% in at least one individual. The observed number of PTVs per individual is consistent with the ~100 loss-of-function variants observed in the 1000 Genomes project ^9^. In contrast, the number of PTV singletons (or observed allele counts less than 10) in ExAC suggests approximately five singletons per individual and only ~0.2 per individual in highly constrained genes ^10,11^. These observations indicate that the majority of PTVs in an individual are common (or common and low frequency) such that they can be assessed via genotyping.

**Figure S1.**
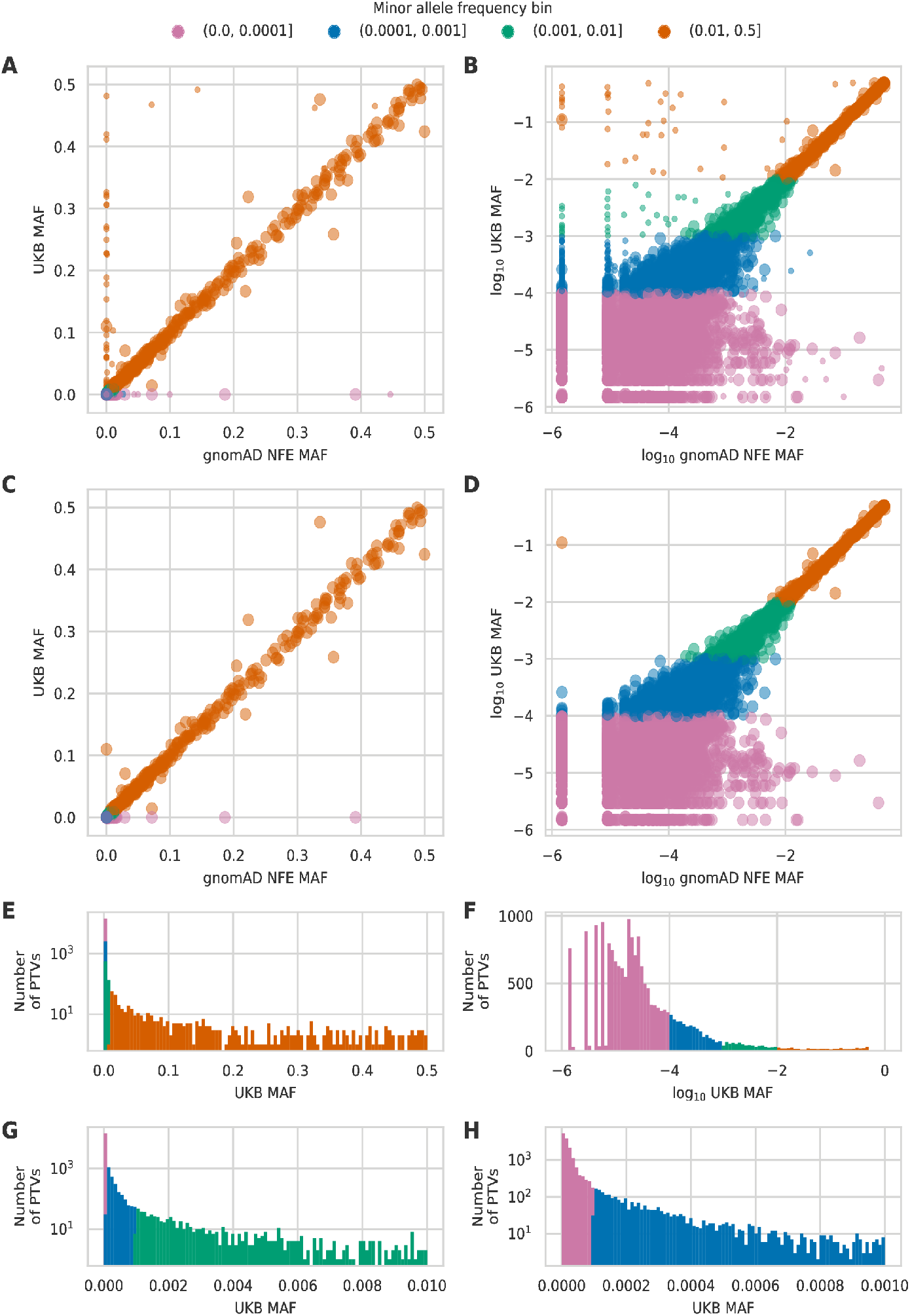
gnomAD allele frequency comparison. (A) PTV allele frequency and (B) log_10_ allele frequency among gnomAD non-Finnish Europeans (NFE) exome dataset and 337,208 UK Biobank (UKB) participants used in this study for all PTVs that could be matched between the two datasets. Small scatter points indicate PTVs that were filtered out for any reason. (C) PTV allele frequency and (D) log_10_ allele frequency among gnomAD non-Finnish Europeans exome dataset and 337,208 UK Biobank participants used in this study for PTVs that passed filtering and could be matched between the two datasets. (E) MAF histogram for all 18,228 polymorphic PTVs that passed filtering. (F) log_10_ MAF histogram for all 18,228 polymorphic PTVs that passed filtering. (G) MAF histogram for 17,765 PTVs with MAF < 0.01. (H) MAF histogram for 17,065 PTVs with MAF < 0.001.

We used computational matching and manual curation based on hospital in-patient record data, self-reported verbal questionnaire data, and cancer and death registry data to define a broad set of medical phenotypes including various cancers, cardiometabolic diseases, and autoimmune diseases (Table S1) ^12^. We then performed association analyses between the 3,724 PTVs with MAF greater than 0.01% and 135 medical phenotypes with at least 2,000 case samples (Figure 1, Figure S2) and stratified the association results into three bins based on PTV MAF greater than 1% (463 PTVs), between 0.1% and 1% (700 PTVs), and between 0.01% and 0.1% (2,561 PTVs) to account for expected differences in the statistical power to detect associations for PTVs with different MAFs (Figure S3). We adjusted the nominal association p-values separately for each MAF bin using the Benjamini-Yekutieli (BY) procedure to correct for multiple hypothesis testing and identified 74 significant associations between PTVs and medical phenotype (BY-adjusted p < 0.05, Figure 2A-C, Table S2).

**Figure 1.**
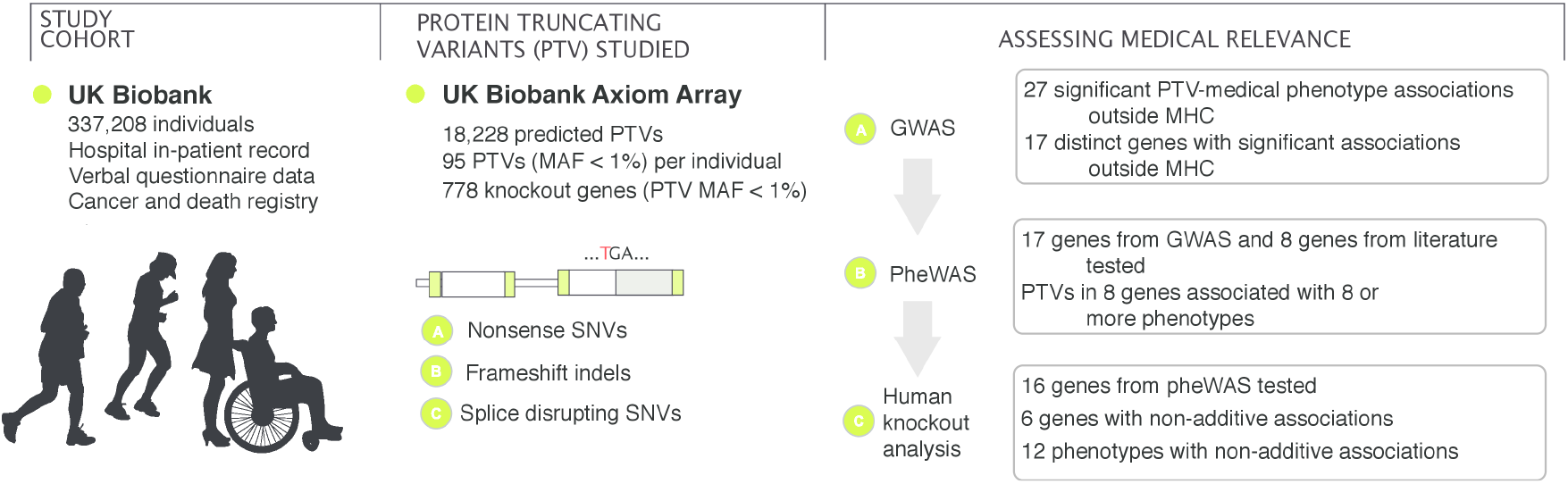
**Schematic overview of the study.** We prepared a dataset of 18,228 protein truncating variants and 135 medical phenotypes from the UK Biobank dataset of 337,208 individuals. From these data, we analyzed the clinical effects of predicted protein-truncating genetic variants.

**Figure S2.**
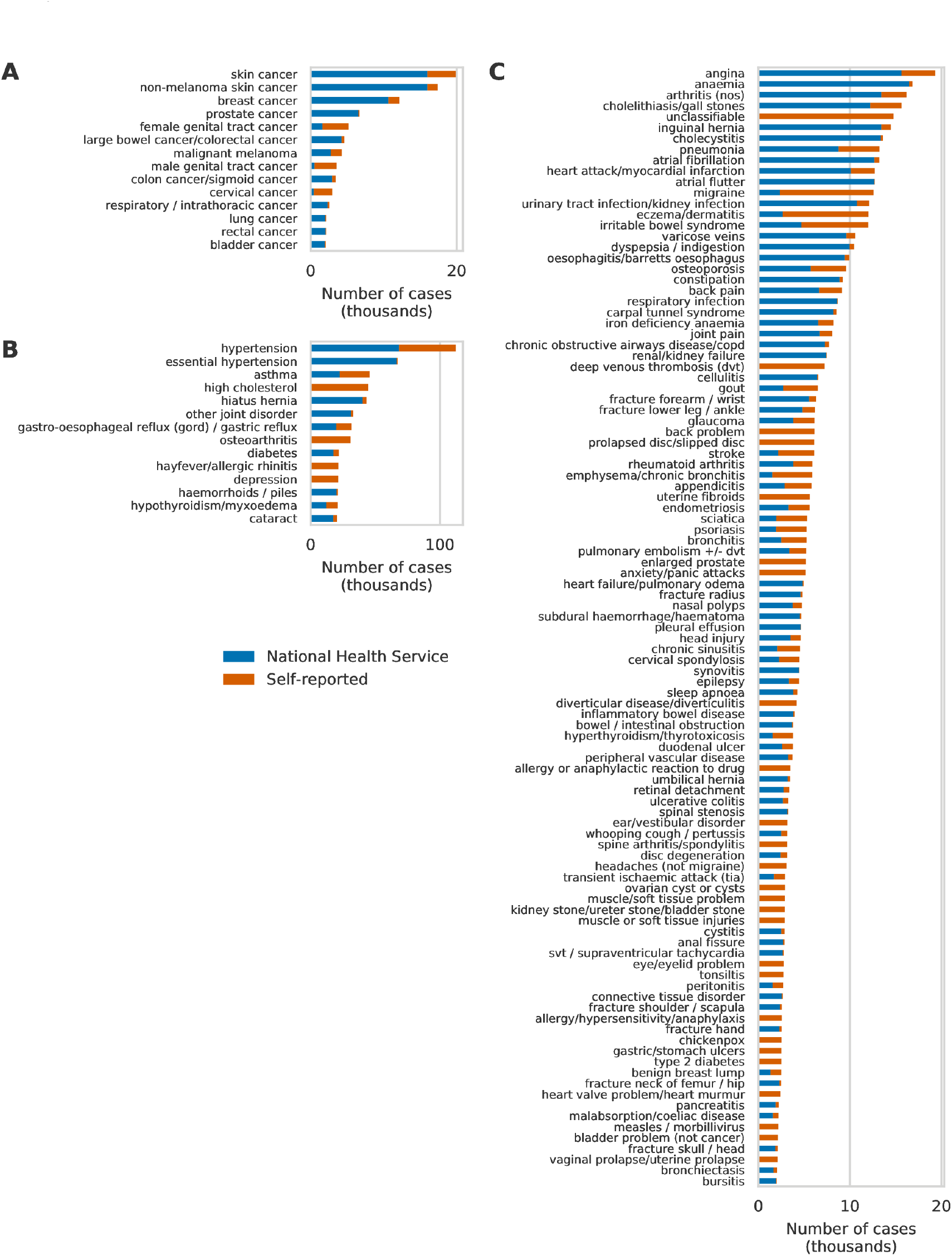
Case numbers for phenotypes with at least 2,000 cases. Number of cases for (A) cancers with more than 2,000 cases, (B) high confidence phenotypes with more than 20,000 cases, and (C) high confidence phenotypes with more than 2,000 cases but less than 20,000 cases. Bars are colored according to the number of cases identified from health records (blue) or questionnaire data (orange).

**Figure S3.**
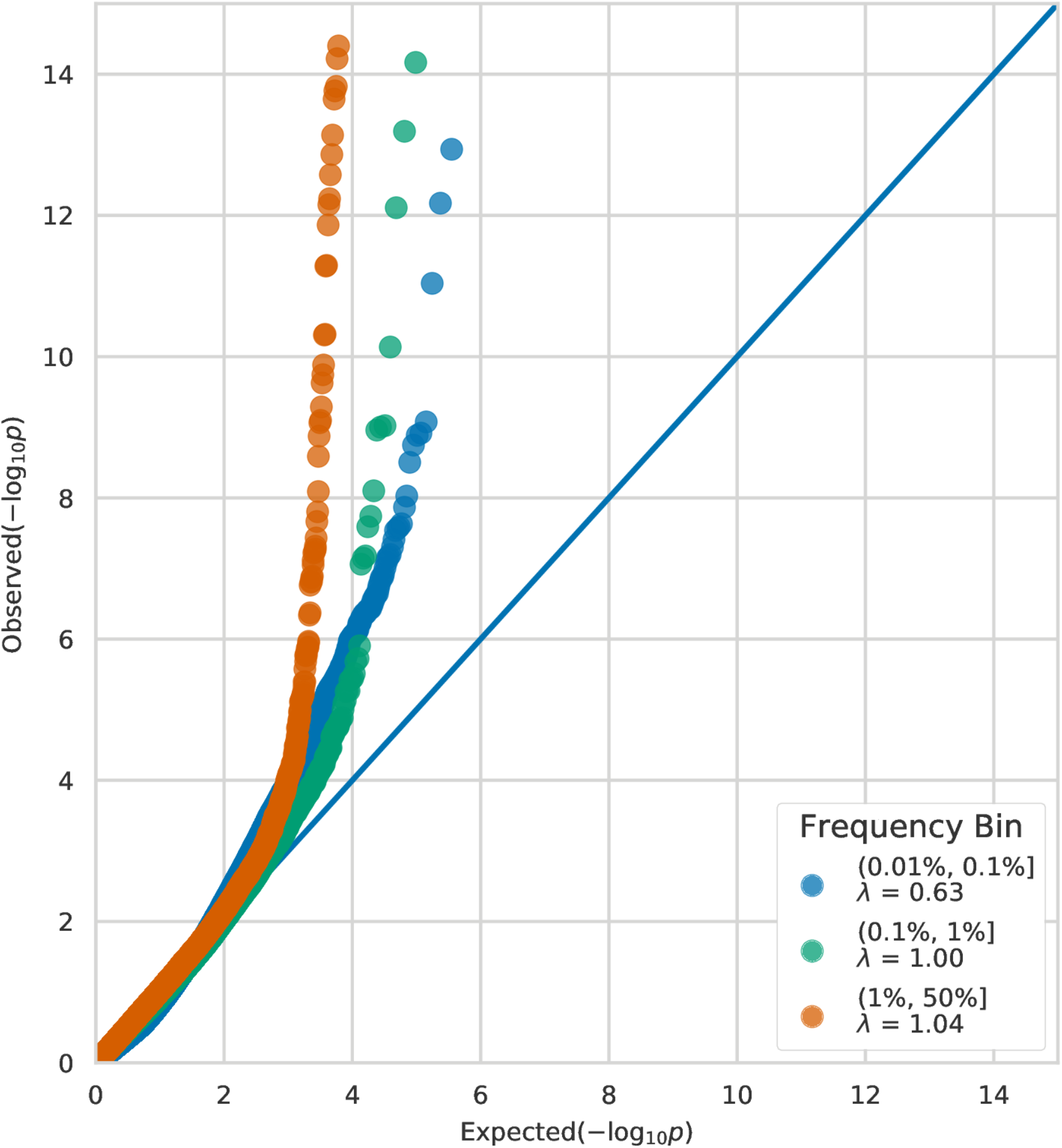
GWAS QQ plots. QQ plots for single variant association analyses for 3,724 PTVs stratified into three minor allele frequency bins: (0.01%, 0.1%], (0.1%, 1%], (1%, 50%]. 26 associations with -log_10_ p-values greater than 14 are not shown.

Among the 74 PTV-phenotype associations we identified, 27 involved PTVs in genes outside of the MHC. We identified five PTVs with seven associations consistent with protective effects (odds ratio [OR]<1, BY-adjusted p<0.05, Figure 2D, Table S2). We found that the rare splice-disrupting PTV rs146597587 in *IL33* is strongly associated with protection against asthma (MAF=0.48%, p=7.6x10^−13^, OR=0.64, 95% CI: 0.57-0.72). This PTV is negatively associated with eosinophil counts (β=-0.21 SD, p=2.5×10^−16^) and has suggestive evidence of an association with asthma (p=1.8x10^−4^, OR=0.47, 95% CI: 13 0.32-0.70)^13^. Our results provide strong evidence in an independent sample that this PTV protects against asthma and suggests that knocking down *IL33* function may be a useful therapeutic approach for asthma. We also identified protective associations for the PTV rs11078928 (MAF=47.1%) in *GSDMB* against asthma (p=6.3x10^−50^, OR=0.90, 95% CI:0.88-0.91) and bronchitis (p=2.6x10^−6^, OR=0.91, 95% CI: 0.87-0.95). *GSDMB* is associated with asthma in humans and induces an asthma phenotype in mouse when overexpressed ^14,15^. We identified additional protective associations between PTVs in *IFIH1* and hypothyroidism (labeled as hypothyroidism/myxoedema) (MAF=1.5%, p=1.7x10^−6^, OR=0.80, 95% CI: 0.73-0.88) and *VKORC1* and hypertension (MAF=25.3%, p=1.4x10^−6^, OR=0.97, 95% CI: 0.96-0.98).

**Figure 2.**
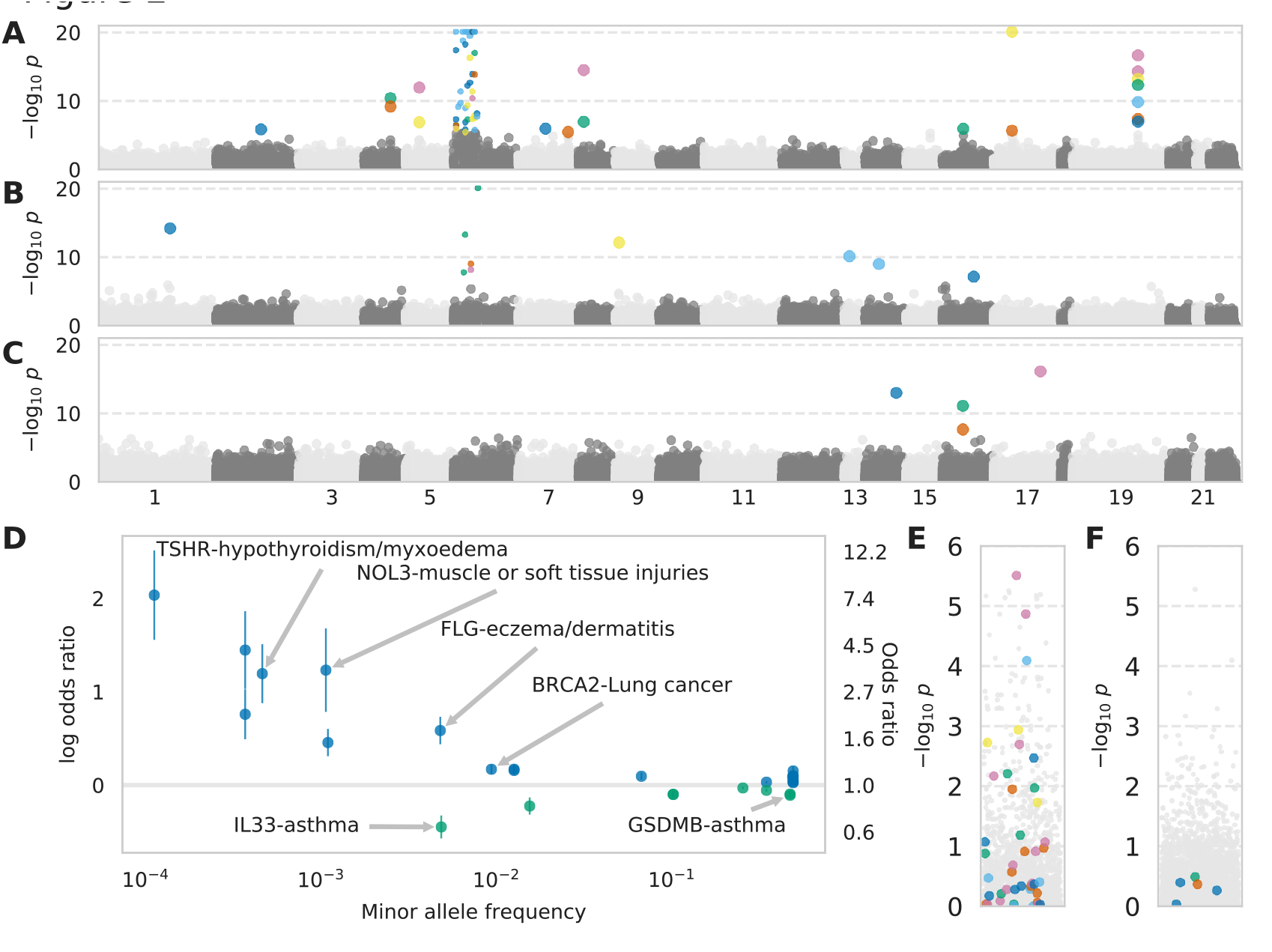
Identification of risk and protective alleles for 135 phenotypes. (A-C) Manhattan plots for all PTVs and all phenotypes stratified by minor allele frequency (A) greater than 1%, (B) between 0.1% and 1%, and (C) between 0.01% and 0.1%. Scatter points are colored according to phenotype. 14 associations with -log_10_ p-values greater than 20 were plotted at 20. PTVs in genes near or in the MHC region have smaller scatter points. (D) Effect size “cascade plot” for all associations outside the MHC with BY-adjusted p < 0.05. Error bars represent 95% confidence intervals. (E-F) Manhattan plots for PTVs in or near the MHC with minor allele frequency (E) greater than 1% and (F) between 0.1% and 1%. The p-values for grey points are the same as in (A) and (B), respectively. The p-values for the color points have been recalculated conditional on HLA alleles.

We also found 20 risk associations for PTVs in 12 genes outside the MHC (Figure 2D, Table S2). We identified clinically relevant PTV-phenotype associations such as *FLG*, whose protein product contributes to the structure of epidermal cells, and eczema/dermatitis (MAF=0.48%, p=6.7x10^−15^, OR=1.80, 95% CI: 1.55-2.08) ^16^ and *TSHR,* thyroid stimulating hormone receptor, and hypothyroidism/myxoedema (MAF=0.046%, p=1.2x10^−13^, OR=3.30, 95% CI: 2.41-4.53) ^17^. We replicated known risk genome-wide association study (GWAS) associations such as *BRCA2* and family history of lung cancer (MAF=0.93%, p=7.3x10^−11^, OR=1.19, 95% CI: 1.13-1.25) ^18^ and rs33966350 in *ENPEP* and hypertension (MAF=1.3%, p=4.8x10^−11^, OR=1.17, 95% CI: 1.12-1.23) ^19^ and identified risk associations between *FANCM*, a member of the same gene family as *BRCA2*, and lung cancer (MAF=0.11%, p=9.7x10^−10^, OR=1.58, 95% CI: 1.36-1.83) as well as *NOL3*, a regulator of apoptosis in muscle cells, and muscle or soft tissue injury (MAF=0.11%, p=6.5x10^−8^, OR=3.43, 95% CI:2.19-5.36) ^20,21^. Even in the context of variants with strong predicted effects such as PTVs, it is critical to evaluate whether the associated variant is causal in the context of neighboring variants. We initially identified an association between the PTV rs34358 in *ANKDD1B* and high cholesterol, although this association disappeared upon conditional analysis with rs17238484, an intronic variant in *HMGCR* known to be associated with cholesterol levels ^22^. Another association between rs34358 and family history of diabetes remained upon conditional analysis with rs17238484 (p=9.1x10^−5^, OR=1.03, 95% CI: 1.02-1.05). Overall we found both PTV-phenotype associations that reflect known biology or disease associations and PTV-phenotype associations that implicate genes in disease.

We identified five significant associations between PTVs and family history phenotypes included in our analysis (Table S2). For two of these associations, the variant associated with the family history phenotype was also associated directly with the phenotype. rs180177132 in *PALB2* was associated with a family history of breast cancer (MAF=0.037%, p=2.5x10^−8^; OR=2.14, 95% CI: 1.64-2.79) as well as breast cancer diagnosis (p=9.0x10^−12^; OR=4.25, 95% CI: 2.80-6.43) and *FUT2* was associated with family history of high blood pressure (MAF=49.1%, p=1.3x10^−7^; OR=1.03, 95% CI: 1.02-1.04), hypertension diagnosis (p=5.7x10^−13^; OR=1.04, 95% CI: 1.03-1.05), and essential hypertension (p=5.2x10^−8^, OR=1.04, 95% CI: 1.02-1.05). We also found that the PTV rs11571833 in *BRCA2* was associated with lung cancer (MAF=0.934%, p=7.3x10^−11^, OR=1.19, 95% CI: 1.13-1.25). These results demonstrate previous approaches for identifying genetic associations using family history information (e.g. ^23^) can be applied even to relatively rare PTVs.

To further characterize the PTV-phenotype associations, we asked whether missense variants with MAF greater than 0.01% in the genes with significant PTV associations were also associated with the same phenotypes. For each of the 27 PTV-phenotype associations in our GWAS, we performed association analyses between the missense variants in that gene and the phenotype that the PTV was associated with and found 23 missense variant-phenotype associations with p<0.001 (Table S2). 13 of these 23 associations remain significant when after a conditional analysis including the PTV genotype as a covariate indicating that a number of genes with PTV associations also contain independent missense associations. For instance, we found two different missense variants in *TSHR* that were both associated with hypothyroidism independent of the PTV association. We also identified independent missense associations for genes and phenotypes such as *ENPEP* and hypertension; *GSDMB* and asthma; *IFIH1* and hypothyroidism; and *PALB2* and lung cancer (Table S2). In total, we found at least one missense association for seven genes implicated in our PTV GWAS providing more evidence that these genes are likely important to the etiology of these conditions.

47 of the 74 significant associations involved PTVs in genes in or near the MHC (Table S2). To investigate whether these associations are caused by linkage between these PTVs and HLA susceptibility alleles, we performed association analyses for each of these PTVs conditional on the presence of each of 344 HLA alleles that were polymorphic among the 337,208 subjects (Table S3). We found that the p-values for all five associations with MAF between 0.1% and 1% were greater than 0.05 for at least one HLA allele (Figure 2E). Similarly, the p-values for 30 of 42 associations with MAF greater than 1% were greater than 0.05 for at least one HLA allele and only three were less than 0.001 (Figure 2F). For instance, we identified an association between rs72841509 in *BTN3A2* and Celiac disease (coded malabsorption/coeliac disease) in our initial GWAS (MAF=0.13, p=1.8x10^−119^, OR=2.33, 95% CI: 2.17-2.50). However, conditioning upon the presence of the well-known Celiac disease risk allele HLA-B8 reduced the p-value of the association between rs72841509 and Celiac disease to p=0.92 ^24^. These results indicate that the majority of the associations identified here for PTVs in MHC genes are likely due to LD with HLA susceptibility alleles and show that it is important to carefully consider the genomic context of associated variants, even for variants with strong predicted effects ^25^.

We next investigated whether we could identify PTV-phenotype associations using imputed genotypes. After filtering (Methods), we identified 546 PTVs outside the MHC with MAF greater than 0.01% among the UK Biobank imputed genotypes. We stratified these PTVs into the same MAF bins as above (0.01%-0.1%, 0.1%-1%, and 1%-50%) and applied the BY adjustment to the association p-values for each bin. We found nine significant associations for imputed PTVs (BY-adjusted p<0.05, Table S2) including rs74315329 in *MYOC* and glaucoma (MAF=0.0012, p=1.8x10^−30^, OR=4.71, 95% CI: 3.61-6.14) ^26^, a well-known risk variant for glaucoma ^27^, and *D2HGDH* and asthma (MAF=0.445, p=1.6x10^−12^, OR=0.95, 95% CI: 0.94-0.96) and hay fever (coded hayfever/allergic rhinitis) (p=8.4x10^−9^, OR=0.94, 95% CI: 0.92-0.96). The *D2HGDH* PTV is in partial LD with an intronic variant rs34290285 in *D2HGDH* (r^2^=0.366, LDlink) that has been associated with asthma and allergic disease^28,29^. We also identified an association between the PTV rs754512 in *MAPT* and Parkinson’s disease (MAF=0.23, p=1.1x10^−6^; OR=0.94, 95% CI: 0.92-0.97) ^30^. This variant is predicted to be a PTV but is in the intron of the canonical *MAPT* transcript and lies on the same haplotype as three *MAPT* missense variants (rs17651549, rs62063786, rs10445337) so conditional analysis could not establish the causal allele. We found associations between a PTV in *RPL3L* and atrial flutter (MAF=0.0021, p=5.0x10^−10^, OR=0.54, 95% CI: 0.44-0.66) and atrial fibrillation (p=2.3x10^−9^, OR=0.55, 95% CI: 0.46-0.67). The missense variant rs140185678 in *RPL3L* is also independently associated with atrial fibrillation (MAF=0.0363, p=5.4x10^−9^, OR=1.21, 95% CI: 1.14-1.30, unpublished data) and atrial flutter (p=1.1x10^−7^, OR=1.20, 95% CI: 1.12-1.28). Overall, we were able to recover a small number of associations using imputed PTVs, indicating that better imputation methods are likely needed in the absence of direct genotyping of PTVs.

To further assess the role of PTVs across medical phenotypes, we performed a phenomewide association analysis (pheWAS) to determine whether PTVs that have been implicated in disease predisposition may impact other diseases or commonly measured traits ^31^. We focused this analysis on PTVs with minor allele frequency greater than 0.01% in the 17 genes with significant associations in our GWAS. In addition to PTVs in the genes identified here, we also investigated PTVs in genes with previously identified protective effects such as: *CARD9, RNF186* and *IL23R* shown to confer protection against Crohn’s disease and/or ulcerative colitis ^2,1^; *ANGPTL4*, *PCSK9*, *LPA*, and *APOC3*shown to confer protection against coronary heart disease ^4,32,33,34,35,36^; and *SCN9A* where homozygous PTV carriers show an inability to experience pain ^37^ (Table S4).

We identified all associations (p<0.01) for PTVs in these 25 genes with a MAF greater than 0.01% and found that PTVs in many of these genes were associated with a broad range of phenotypes (Table S2, Figure S4). PTVs in eight of the 25 genes were associated with eight or more phenotypes. We observed associations between the viral receptor *IFIH1* and 10 phenotypes including protective effects against hypothyroidism, hypertension, gastric reflux, and psoriasis (Figure 3, Table S4). Despite minor allele frequencies ranging from 0.02% to 1.5%, three of these associations were observed for more than one *IFIH1* PTV. PTVs in *IFIH1* were also associated with increased risk for ulcerative colitis, inflammatory bowel disease, and endometriosis. We identified new protective effects for *IL33* for hay fever (coded hayfever/allergic rhinitis), nasal polyps, and angina as well as weak risk effects for bowel/intestinal obstruction and shoulder/scapula fracture (Figure S4). Overall, these results demonstrate that PTVs can have pleiotropic effects across diverse phenotypes and that PTVs in the same gene can both protect against and increase risk for different diseases.

**Figure 3.**
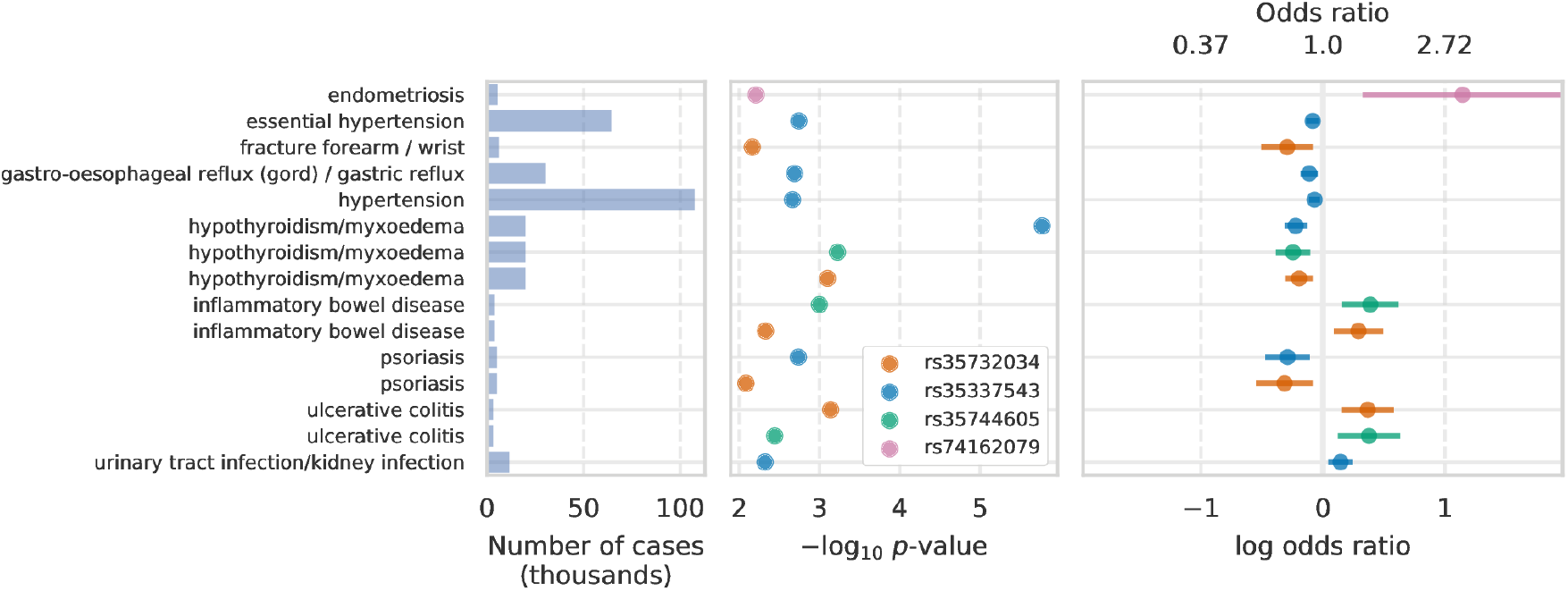
PheWAS for *IFIH1*. Phenome-wide associations (p<0.01) for four PTVs in *IFIH1* with minor allele frequency greater than 0.01%. The left panel shows the number of cases per phenotype in thousands. The middle panels shows the logistic regression -log_10_ p-value. The right panel shows the estimated odds ratios and 95% confidence intervals.

**Figure S4.**
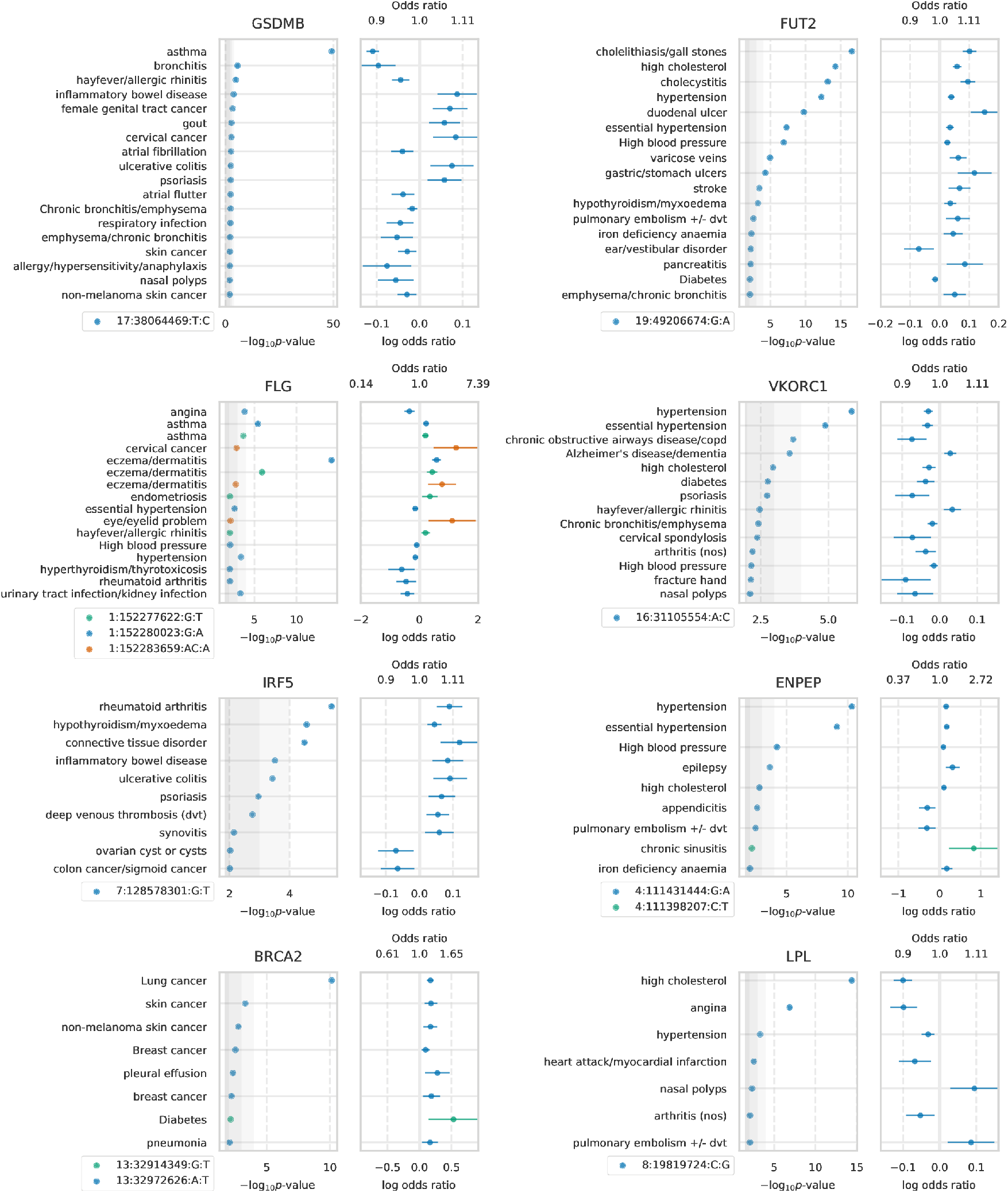
PheWAS results. -log_10_ p-values and odds ratios for associations (p<0.01) for PTVs in 21 genes and 135 medical phenotypes. The gene for each plot is indicated above the p-value panel. *IFIH1* is plotted in Figure 3.

Homozygous carriers of PTVs, referred to as homozygous knockouts (KOs), may have dramatically altered medical outcomes compared to carriers with only one PTV 38 (heterozygous KOs)^38^. Genetic association analyses typically assume that genetic effects are additive; that is, the log OR of a homozygote is expected to be twice the log OR of a heterozygote. Given the large difference between having one functional copy and no functional copies of a gene, however, we expect that homozygote KOs may have non-additive effects that are stronger or weaker than would be predicted given the effect size for heterozygote KOs. To assess whether any of the genes whether any of the 17 genes with significant associations in our GWAS or the eight genes with published protective effects (Table S4) have evidence for non-additive effects on medical phenotypes, we estimated the KO status in each subject for each of these 25 genes. Subjects with one PTV in a gene were considered heterozygote KOs for that gene and subjects with two or more PTVs were considered homozygote KOs. In total, 16 of the 25 genes had at least one predicted homozygous KO carrier. We fit additive and non-additive models to test for associations between KO status for these 16 and 206 medical phenotypes (minimum 1,000 cases, Figure S5) and found 13 associations (6 distinct genes, 12 distinct phenotypes) with potential non-additive effects (Figure S6, Table S5, Methods).

**Figure S5.**
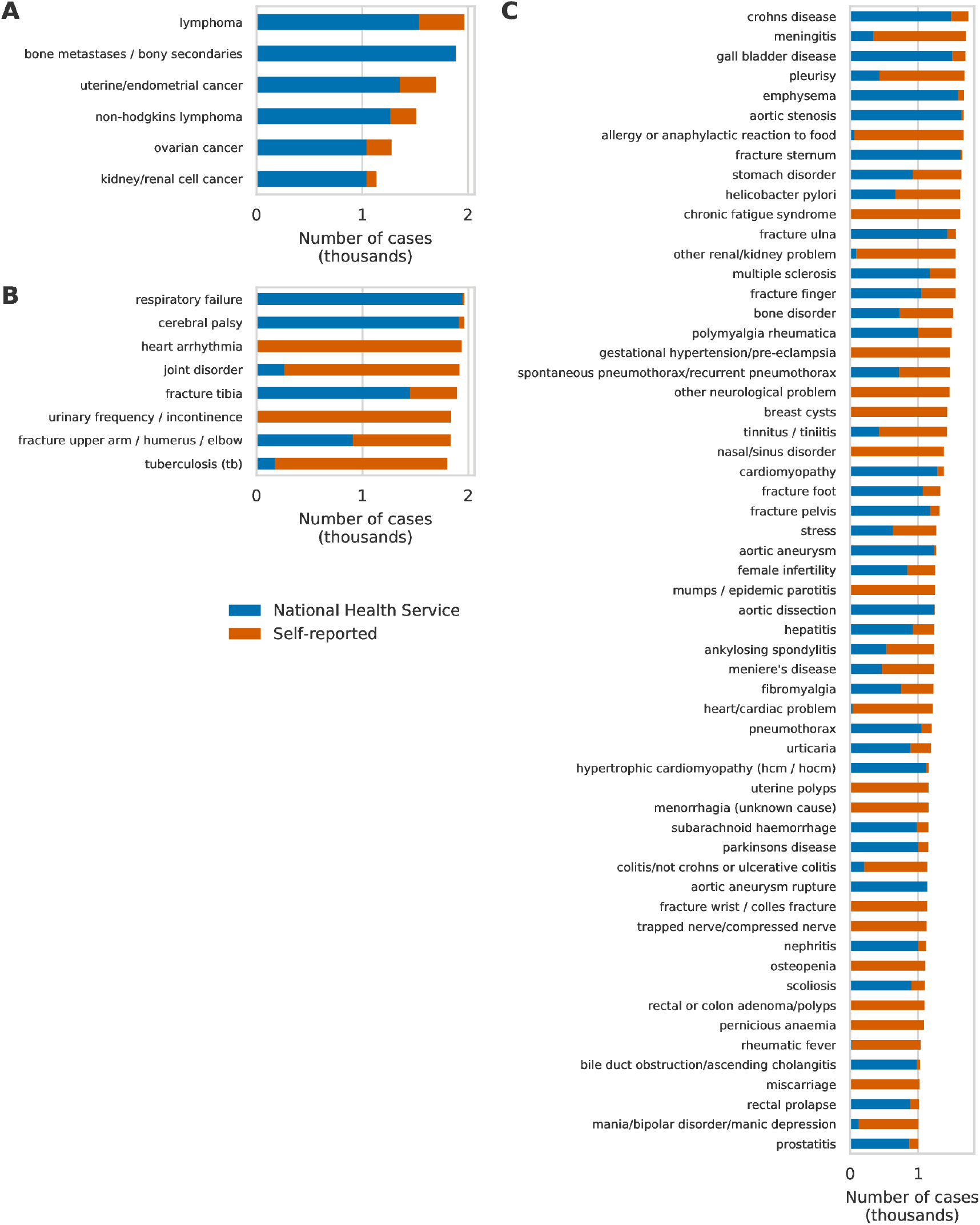
Case numbers for phenotypes with between 1,000 and 2,000 cases. Number of cases for (A) cancers with more than 1,000 cases but less than than 2,000 cases, (B) high confidence phenotypes with more than 1,750 cases but less than 2,000 cases, and (C) high confidence phenotypes with more than 1,000 cases but less than 1,750 cases. Bars are colored according to the number of cases identified from health records (blue) or questionnaire data (orange).

**Figure S6.**
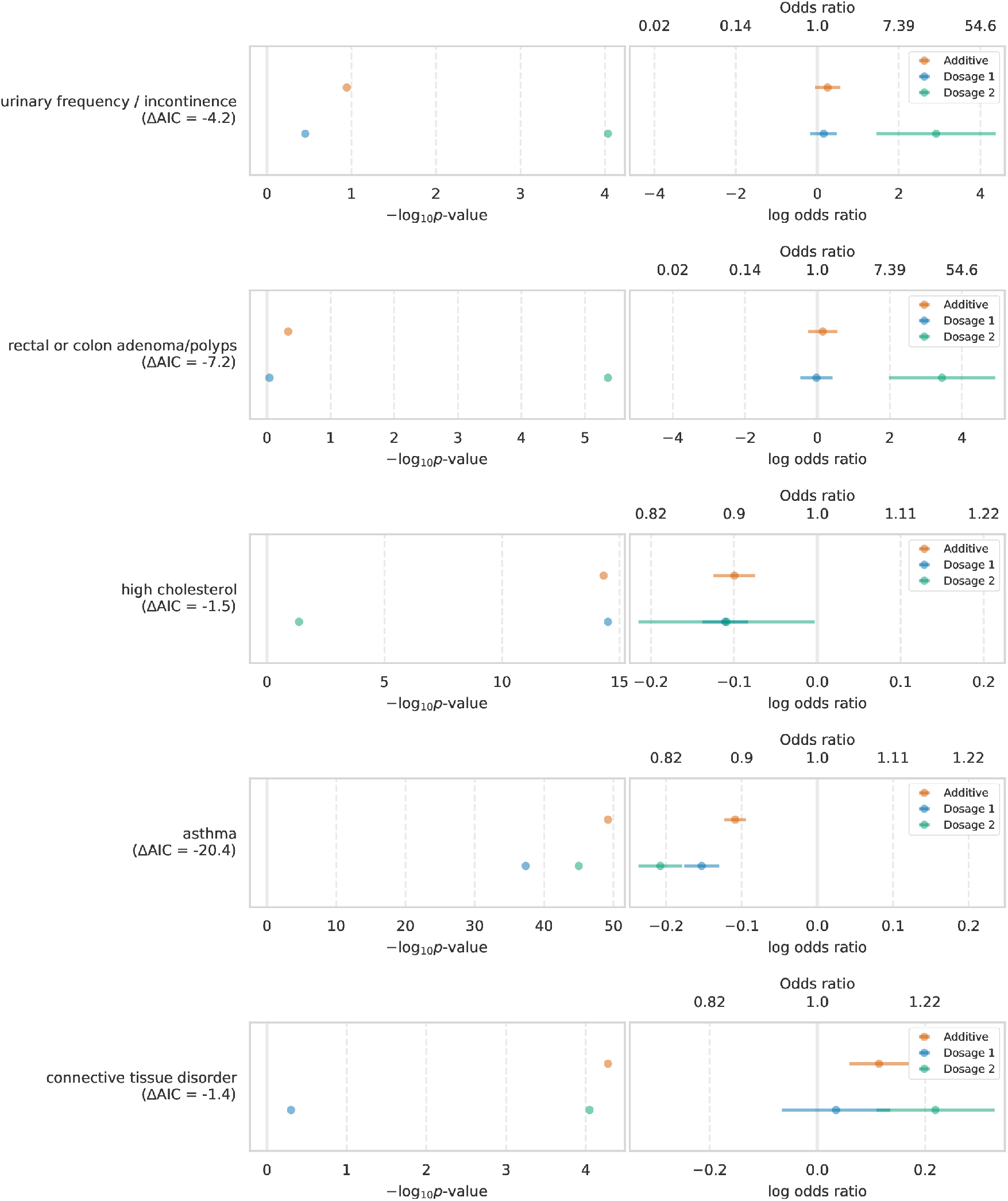
Non-additive associations. The left panel shows the -log_10_ p-values for (blue) additive genetic model and (green, orange) non-additive genetic model. The right panel shows the estimated odds ratio and 95% confidence interval for (blue) additive genetic model and (green, orange) non-additive genetic model. The gene-phenotype association is labeled on the left with the difference in AIC between the additive and non-additive models. A more negative difference in AIC favors the non-additive model. *FUT2* is plotted in Figure 4.

We identified 87,176 predicted homozygous KOs for *FUT2* caused by a common PTV rs601338 with MAF 49.1% and identified non-additive risk associations between *FUT2* KO status and eight phenotypes including hypertension and mumps (Figure 4, Table S5). FUT2 regulates the expression of the H antigen on the gastrointestinal mucosa and genetic variation in *FUT2* is associated with Crohn’s disease ^39,40^, psoriasis ^41^, plasma vitamin B12 levels ^42,43^, levels of two tumor biomarkers ^44,45^, and urine fucose levels ^46^. Under a non-additive model, the ORs for heterozygous *FUT2* KOs are all nearly one while *FUT2* homozygous KOs have ORs ranging from 1.05 (95% CI: 1.03-1.07) to 1.51 (95% CI: 1.29-1.77). Given the frequency of the rs601338 PTV, our results indicate that FUT2 function may play an important role in a wide range of phenotypes.

**Figure 4.**
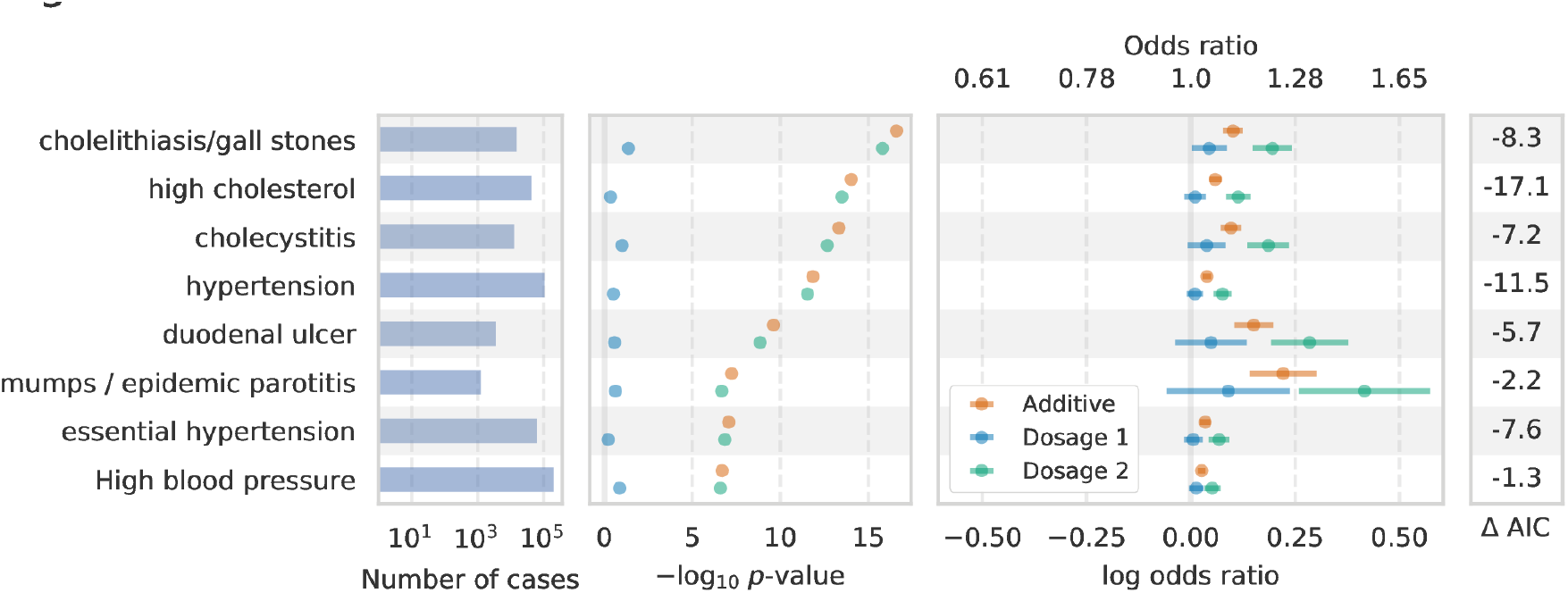
Non-additive associations for *FUT2*. Association results under additive and non-additive models for predicted *FUT2* heterozygous or homozygous knockouts (KOs) with a difference between nonadditive model AIC and additive model AIC less than -1. The left panel shows the number of cases per phenotype. The middle-left panel shows the -log_10_ p-value for the KO association analysis. The middle-right panel shows the estimated log odds ratios and 95% confidence intervals under an additive model (orange) and under a non-additive model for heterozygote KOs (blue) and homozygote KOs (green).

We also found evidence that the association between *GSDMB* KO and asthma described in our GWAS analysis above is non-additive (Figure S6, Table S5). In total, we identified 168,025 heterozygous KOs and 74,534 homozygous KOs for *GSDMB*. Under an additive model, *GSDMB* heterozygote KOs are predicted to have a decreased risk for asthma with OR=0.90 (p=5.9x10^−50;^ 95% CI: 0.88-0.91). Under a non-additive model, however, *GSDMB* heterozygote KOs are predicted to have OR=0.86 (p=4.3x10^−38^; 95% CI: 0.84-0.88) while *GSDMB* homozygote KO offers only modestly higher protection (p=9.7x10^−46^, OR=0.81, 95% CI: 0.79-0.84). Variants that increase expression of *GSDMB* in humans 47 are associated with asthma risk,^47^ and increased *GSDMB* expression causes an asthma phenotype in mice ^48^. Our results suggest that knocking out just one copy of *GSDMB* provides most of the effect on asthma risk. Overall, we identified non-additive PTV associations for six of 16 genes tested demonstrating that the effect of PTVs on disease risk can be complex.

## Discussion

Assessing the medical relevance of protein-truncating variants is critical for prioritizing putative drug targets and clinical interpretation. We systematically characterized the association of PTVs, a class of variants with functional consequences likely to be consistent with inhibition, with medical phenotypes using data from the UK Biobank study. We estimated the effects of PTVs across 135 phenotypes and identified 27 associations between PTVs in 17 genes and 20 different phenotypes. We found four associations for PTVs with minor allele frequency less than 0.1% indicating that more subjects or case/control studies design may be needed to test for associations between ultra-rare PTVs and relatively low prevalence diseases that are not well-represented in biobank datasets. We performed 25 phenome-wide association analyses for the genes implicated by GWAS in this study plus eight genes curated from the literature (Table S4) and identified eight genes that were associated with eight or more phenotypes (p < 0.01). 6 of these 25 genes showed evidence for non-additive associations across several phenotypes including non-additive associations between *GSDMB* and asthma and *FUT2* and eight phenotypes including hypertension and cholesterol.

The genetic associations reported here directly link gene function to disease etiology and provide attractive targets for drug discovery. Naturally occurring human knockouts that protect against disease provide *in vivo* validation of safety and efficacy and may be relatively simple to target with drugs. Protective associations between PTVs in *IL33* and asthma; *GSDMB* and asthma; and *IFIH1* and hypothyroidism represent particularly attractive drug targets while risk associations between PTVs in *FANCM* and lung cancer and *NOL3* and muscle injuries implicate these genes as important to the development of these conditions. Our results illustrate the value of deep population-scale health and genomic datasets for prioritizing genetic variants and genes with translational potential.

## Author Contributions

M.A.R. conceived and designed the study. C.D., Y.T., G.M., A.L., and M.A.R. designed and carried out the statistical and computational analyses. C.C. optimized and implemented computational methods. C.D., Y.T., G.M., A.L., and M.J.D. carried out quality control of the data. Browser features in Global Biobank Engine were led and developed by G.M. and M.A.R, with assistance from A.L., Y.T., and C.D. M.A.R. supervised all aspects of the study. M.J.D. and C.D.B. provided analysis and commented on the manuscript. The manuscript was written by C.D., Y.T., and M.A.R.

## Acknowledgements

This research has been conducted using the UK Biobank resource.We thank all the participants in the UK Biobank study. We would like to thank Stefan Stender for suggesting that the association between the PTV rs34358 and high cholesterol might be due to the LDL GWAS variant rs17238484 in *HMGCR* (https://gitter.im/UK-Biobank/Lobby). M.A.R. is supported by Stanford University and a National Institute of Health center for Multi - and Trans-ethnic Mapping of Mendelian and Complex Diseases grant (5U01 HG009080). C.D. is supported by a postdoctoral fellowship from the Stanford Center for Computational, Evolutionary, and Human Genomics. Y .T. is supported by Funai Overseas Scholarship from Funai Foundation for Information Technology and the Stanford University Biomedical Informatics Training Program. A.L. and G.M. are supported by NIH BD2K grant number T32 LM 012409. The primary and processed data used to generate the analyses presented here are available in the UK Biobank access management system (https://amsportal.ukbiobank.ac.uk/) for application 24983, “Generating effective therapeutic hypotheses from genomic and hospital linkage data” (http://www.ukbiobank.ac.uk/wp-content/uploads/2017/06/24983-Dr-Manuel-Rivas.pdf), and the results are displayed in the Global Biobank Engine (https://biobankengine.stanford.edu). We would like to thank the Customer Solutions Team from Paradigm4 who helped us implement efficient databases for queries and application of inference methods to the data.

## Supplementary Materials

### Materials and Methods

#### Quality Control of Genotype Data

We used genotype data from UK Biobank dataset release version 2 for all aspects of the study except the imputed PTV GWAS ^49,50^. To minimize the impact of cofounders and unreliable observations, we used a subset of individuals that satisfied all of the following criteria: (1) self-reported white British ancestry, (2) used to compute principal components, (3) not marked as outliers for heterozygosity and missing rates, (4) do not show putative sex chromosome aneuploidy, and (5) have at most 10 putative third-degree relatives. We removed 151,169 individuals that did not meet these criteria. For the rest of 337,208 individuals, we used PLINK v1.90b4.4 ^51^ to compute the following statistics for each variant: (a) genotyping missingness rate, (b) p-values of Hardy-Weinberg test, and (c) allele frequencies.

#### Protein-Truncating Variant Annotation

We annotated 784,257 autosomal variants extracted from the mapping bim files provided by the UK Biobank using VEP version 87 and the LOFTEE plugin (https://github.com/konradjk/loftee) and identified 27,057 putative PTVs ^52^. We first removed 8,118 PTVs specific to the UK BiLEVE Axiom Array or with missingness greater than 1% among the subjects genotyped on the UK Biobank Axiom Array. Despite a missingness rate of 28% on the Axium Biobank Array, we kept rs141992399 (*CARD9*) in the analysis. We removed 11 variants with cluster plots that indicated unreliable genotypes. We removed Affx-89018997 because the REF/ALT annotation caused problems with analysis software.

We next matched our PTVs to PTVs annotated in gnomAD (gnomad.exomes.r2.0.1.sites.vcf.gz) based on genomic position, reference, and alternate alleles and compared the allele frequencies in the UKB and gnomAD by (1) performing a Fisher’s exact test using the minor allele counts from the 337,208 UKB subjects and the minor allele counts from gnomAD and (2) calculating the log odds ratio of observing the minor allele in the UKB versus gnomAD. We stratified our PTVs by minor allele frequency into the following three bins: (0.01%, 0.1%], (0.1%, 1%], (1%, 50%]. For bin (0.01%, 0.1%], we removed PTVs with Fisher p < 1e-7 and an absolute log odds ratio greater than 3. For bin (0.1%, 1%], we removed PTVs with Fisher p < 1e-7 and an absolute log odds ratio greater than 2. For bin (1%, 100%], we removed PTVs with Fisher p < 1e-7 and an absolute log odds ratio greater than 1 (Figure S1). In total, 123 variants were removed in this step.

There were 134 variants with MAF greater than 0.1% that we did match to the gnomAD exome data. We manually reviewed these variants on the gnomAD browser to determine whether they were likely to accurately type a PTV in gnomAD. In cases where the PTV was present on the gnomAD browser but was not included in the exome data, we kept the PTV in our analysis. In cases where the UKB array likely typed a non-PTV or there was no variant present on the browser, we removed the PTV from our analysis. In total, 79/134 variants were removed during in this step. 18,726 PTVs remained after filtering of which 18,228 were polymorphic. We focused on these 18,228 PTVs for subsequent analyses.

We considered variants on chr6:25477797-36448354 as in or near the MHC for all analyses. We use the hg19 human genome reference throughout.

### Cancer Phenotype Definitions

We used the following procedure to define cases and controls for cancer GWAS. Individual level ICD-10 codes from the UK Cancer Register (http://biobank.ctsu.ox.ac.uk/crystal/label.cgi?id=100092), Data-Field 40006 (http://biobank.ctsu.ox.ac.uk/crystal/field.cgi?id=40006), and the National Health Service (http://biobank.ctsu.ox.ac.uk/crystal/label.cgi?id=2022), Data-Field 41202 (http://biobank.ctsu.ox.ac.uk/crystal/field.cgi?id=41202), in the UK Biobank were mapped to the self-reported cancer codes, Data-Field 20001 (http://biobank.ctsu.ox.ac.uk/crystal/field.cgi?id=20001). The mapping was performed via manual curation of ICD-10 codes for each of the self-reported cancer codes. UKB field codes for self-reported cancer were created with a tree structure such that more specific cancer subtypes (e.g. “malignant melanoma”) are nested under more general categories (“skin cancer”). This tree structure was preserved in the field code to ICD-10 mapping. For example, the self-reported phenotype of “lip cancer” was mapped to its field code, 1010, and the ICD-10 codes for “malignant neoplasm of lip”, C00 and C000-C009. After this mapping, individuals with an affirmative entry in one or more of the phenotype collections (self-reported cancer, cancer registry, and the NHS) were included in the case cohort for the GWAS. No secondary neoplasms were included in the cancer phenotype mappings.

### High Confidence Phenotype Definitions

We used the following procedure to define cases and controls for non-cancer phenotypes (referred to as “high confidence” phenotypes). ICD-10 codes (Data-Field 41202), were grouped with self-reported non-cancer illness codes (Data-Field 20002) that were closely related. This was done by first creating a computationally generated candidate list of closely related ICD-10 codes and self-reported non-cancer illness codes, then manually curating the matches. The computational mapping was performed by calculating the token set ratio between the ICD-10 code description and the self-reported illness code description using the FuzzyWuzzy python package. The high scoring ICD-10 matches for each self-reported illness were then manually curated to ensure high confidence mappings. Manual curation was required to validate the matches because fuzzy string matching may return words that are similar in spelling but not in meaning. For example, to create a hypertension cohort the code description from Data-Field 20002 (“Hypertension”) was mapped to all ICD-10 code descriptions and all closely related codes were returned (“I10: Essential (primary) hypertension” and “I95: Hypotension”). After manual curation code I10 would be kept and code I95 would be discarded.

### Family History Phenotype Definitions

We used data from Category 100034 (Family history - Touchscreen - UK Biobank Assessment Centre) to define “cases” and controls for family history phenotypes. This category contains data from the touchscreen questionnaire on questions related to family size, sibling order, family medical history (of parents and siblings), and age of parents (age of death if died). We focused on Data Coding 20107: Illness of father and 20110: Illness of mother.

### Genome-Wide Association Analyses

We performed genome-wide logistic regression association analysis with Firth-fallback using PLINK v2.00a(17 July 2017). Firth-fallback is a hybrid algorithm which normally uses the logistic regression code described in ^53^, but switches to a port of logistf() (https://cran.r-project.org/web/packages/logistf/index.html) in two cases: (1) one of the cells in the 2x2 allele count by case/control status contingency table is empty (2) logistic regression was attempted since all the contingency table cells were nonzero, but it failed to converge within the usual number of steps. We used the following covariates in our analysis: age, sex, array type, and the first four principal components, where array type is a binary variable that represents whether an individual was genotyped with UK Biobank Axiom Array or UK BiLEVE Axiom Array. For variants that were specific to one array, we did not use array as a covariate. We stratified GWAS p-values from PLINK into three minor allele frequency bins: 0.01%-0.1% (2,562 PTVs), 0.1%-1% (700 PTVs), and >1% (463 PTVs). We corrected p-values separately for each bin using the Benjamin-Yekutieli approach implemented in R’s p.adjust ^54^.

For the missense variant GWAS, we identified missense variants with MAF > 0.01% in each of the 17 non-MHC genes that had a significant PTV from the PTV GWAS. All genes except for *IRF5* had at least one missense variant. We then performed associations analyses as described above for the missense variants from each gene and the phenotypes that PTVs in that gene were associated with. We considered significant any missense-phenotype associations with nominal p<0.001. We repeated the association analyses using the PTV genotype as a covariate to evaluate whether the association signals were independent for significant missense variants.

### HLA Conditional Analysis

We performed conditional association analyses for 47 of the 74 significant associations from our GWAS for PTVs in genes in or near the MHC using the HLA alleles provided by the UK Biobank (ukb_hla_v2.txt). For each PTV-phenotype association, we re-ran the association analysis using each of the 344 HLA alleles polymorphic in the 337,208 subjects used here as a covariate in turn. We then identified which HLA allele, when used as a covariate, corresponded to the largest p-value for the additive genetic effect. These results are reported in Table S3.

### ANKDD1B Conditional Analysis

In our initial GWAS, we found associations between the PTV rs34358 in *ANKDD1B* and family history of diabetes and high cholesterol. Since *ANKDD1B* is near *HMGCR*, we performed a conditional association analysis between rs34358 and family history of diabetes and high cholesterol using the imputed genotypes for rs17238484, an intronic variant in *HMGCR* associated with cholesterol levels ^22^, as covariates. We found that conditioning on rs17238484 made the association between rs34358 and high cholesterol insignificant (p=0.052) but that the association between rs34358 and family history was only slightly reduced from p=1.5x10^−7^ to p=9.1x10^−5^. We therefore decided to include this association in Table S2.

### Imputed PTVs GWAS

We identified 962 PTVs among the UK Biobank imputed genotypes that were not multiallelic, had MAF greater than 0.01%, and were not already included in our study by comparing the chromosomal coordinates and reference and alternate alleles of PTVs annotated in gnomAD to the UK Biobank positions and alleles for the UK Biobank data. We only considered PTVs in the HRC site list version 1.1 (http://www.haplotype-reference-consortium.org/site). We removed 408 imputed PTVs that had an imputation score less than 0.8, missingness greater than 1%, or whose MAF differed substantially from the non-Finnish European MAF in gnomAD. We removed eight more imputed PTVs that were in genes near the MHC. In total we were left with 546 imputed PTVs that we stratified into the following MAF bins: 0.01%-0.1% (247 PTVs), 0.1%-1% (153 PTVs), and >1% (146 PTVs). We corrected p-values separately for each bin using the Benjamin-Yekutieli approach implemented in R’s p.adjust ^54^. We assessed linkage disequlibrium between imputed PTVs and other variants using LDmatrix in LDlink ^55^.

For the missense variant rs140185678 (MAF=0.0363) in *RPL3L,* we ran GWAS as described above and found that the variant was associated with associated with atrial fibrillation (p=5.4x10^−9^, OR=1.21, 95% CI: 1.14-1.30) and atrial flutter (p=1.1x10^−7^, OR=1.20, 95% CI: 1.12-1.28). We re-ran this analysis using the genotype of the *RPL3L* PTV rs140192228 as a covariate and found that the associations between rs140185678 and atrial fibrillation (p=4.3x10^−9^, OR=1.21, 95% CI: 1.14-1.29) and atrial flutter (p=8.4x10^−8^, OR=1.20, 95% CI: 1.12-1.28) were still significant. The PTV was also significant under these models for atrial fibrillation (p=1.1x10^−5^, OR=0.85, 95% CI: 0.79-0.91) and atrial flutter (p=2.1x10^−6^, OR=0.84, 95% CI: 0.78-0.90).

### Phenome-Wide Association Analyses

We performed phenome-wide association analyses (pheWAS) on the 17 genes with at least one significant association in our GWAS as well as 8 genes reported to have protective genetic associations: *CARD9*, *RNF186*, *IL23R*, *ANGPTL4*, *PCSK9*, *LPA*, *APOC3*, and *SCN9A* (Table S4). We identified associations between PTVs in these genes with MAF greater than 0.01% and 135 medical phenotypes (p < 0.01, Figure S4). Four genes (*ANGPTL4*, *IL23R*, *PCSK9*, and *APOC3*) did not have any associations with p<0.01 in the pheWAS.

### Knockout Status

We estimated PTV knockout carrier status for each individual by summing the total number of PTVs present in an individual for each gene that had at least one PTV. If a PTV was predicted to effect more than one gene, we counted that PTV for each gene. If an individual was predicted to carrier more than 2 PTVs in a given gene, we set his or her count to two. We thus obtained carrier statuses for each gene in each subject that ranged from no KO, heterozygous KO, or homozygous KO. For all 18,228 predicted PTVs, we found 262 PTVs per subject on average and 1,173 genes with at least one putative KO. If we restrict to only high confidence (HC) PTVs, we observe 174 PTVs per subject on average and 995 genes with at least one putative KO. If we restrict to PTVs with MAF less than 1%, we observe 95 PTVs per subject on average and 778 genes with at least one putative KO.

### Additivity Analyses

To test for departures from additivity, we tested for associations between PTV carrier status and phenotype status for 16 of the 25 genes used in the pheWAS analysis that had at least one homozygote knockout and 206 phenotypes with at least 1,000 cases. For each gene and phenotype, we fit two models using the glm function in R (family=“binomial”). For the additive model, we provided PTV carrier status as a numeric variable, and for the non-additive model, we provided PTV carrier status as a factor. We included age, sex, genotyping array, and the first four principal components as covariates for both models. To identify gene-phenotype associations with suspected departures from additivity, we identified genes and phenotypes where either the additive p-value or homozygote KO p-value was less than 10^−4^ and the difference between the non-additive model AIC and additive model AIC was less than -1.

### Data Availability

The UK Biobank data is available through the UK Biobank (http://www.ukbiobank.ac.uk/). We will make analysis scripts and notebooks available on Github at publication. GWAS results can be browsed on the Global Biobank Engine (biobankengine.stanford.edu).

### URLs

LDlink, https://analysistools.nci.nih.gov/LDlink/; gnomAD browser, http://gnomad.broadinstitute.org/; UK Biobank, http://www.ukbiobank.ac.uk/.

## Supplemental Tables

**Table S1. Medical Phenotypes.** 206 medical phenotypes used in this study. Category indicates whether the phenotype was derived from family history questionnaire information (FH) or from diagnosis of a cancer (CA) or other disease (HC). See Methods and Figures S2,5 for more information.

**Table S2. Significant GWAS and pheWAS associations.** Significant associations from PTV and missense GWASs and pheWAS analysis. Variant IDs are from the UK Biobank data release. “gwas_protective” and “gwas_risk” tabs contain significant PTV associations (BY-adjusted p<0.05) for genes outside the MHC. “imputed” tab contains significant associations (BY-adjusted p<0.05) for imputed PTVs. “missense” tab contains significant (p<0.001) missense associations. “phewas” tab contains significant (p<0.01) pheWAS associations.

**Table S3. HLA conditional analysis.** p-values for genetic effects for PTVs in MHC genes from our initial GWAS (“P_variant_gwas”) and from an analysis conditional on the HLA allele in the “HLA_subtype” column (“P_variant_conditional”). “P_subtype_conditional” contains the p-value for the association between the given HLA subtype and phenotype in the conditional analysis.

**Table S4. Genes with known protective associations.** We identified eight genes that did not have significant associations in our single-variant analysis (BY-adjusted p<0.05) but have been reported in the literature to protect against different diseases. We included these genes in our pheWAS and additivity analyses.

**Table S5. Non-additive associations.** Results from fitting additive and non-additive models of association for PTV carrier status (no PTVs, heterozygous knockout, or homozygous knockout) in 25 genes against 206 medical phenotypes with at least 1,000 cases. Columns that begin with “dosage1” and “dosage2” correspond to results for the non-additive model while columns that begin with “additive” correspond to the additive model. “aic_diff” is the AIC of the additive model subtracted from the AIC of the non-additive model.

